# Environmental DNA degradation simulation from water temperature and DNA fragment length: A meta-analysis approach

**DOI:** 10.1101/2020.10.30.361980

**Authors:** Tatsuya Saito, Hideyuki Doi

## Abstract

Environmental DNA (eDNA) analysis can detect aquatic organisms, including rare and endangered species, in a variety of habitats. The degradation of eDNA concentration is important to investigate their distribution and has also been experimentally evaluated. It is important to integrate these data to synthesize eDNA degradation in various environments. We collected the eDNA degradation rates and related factors, especially water temperature and fragment lengths of the measured DNA from 28 studies. Our results suggest that water temperature and fragment length are significantly related to the eDNA degradation rate. From the 95% quantile model simulation, we predicted the maximum eDNA degradation rate in various combinations of water temperature and fragment length. Predicting eDNA degradation could be important for evaluating species distribution and inducing innovation of eDNA methods, especially for rare and endangered species with lower DNA concentrations.

## Introduction

Environmental DNA (eDNA) methods are innovative methods developed for monitoring macroorganisms, especially aquatic species (Ficetola et al., 2008; Takahara et al., 2012; Minamoto et al., 2012; Taberlet et al., 2012; Ushio et al., 2018; Tsuji et al., 2019; Kakuda et al., 2019). The eDNA method is used to investigate species distribution, so it is less invasive to the environment and organisms, and is especially useful for rare and endangered species, which generally have low tolerance to sampling disturbance. Consequently, eDNA methods have been used to detect rare and endangered species in various taxa, such as fish, salamander, and aquatic insects (Fukumoto et al. 2015; Sigsgaard et al., 2015; Pfleger et al. 2016; Doi et al., 2017; Sakata et al., 2017).

eDNA, which comprises DNA fragments released by organisms into environments such as water or soil, is thought to be derived from mixtures of feces (Martellini et al., 2005), skin cells (Ficetola et al., 2008), mucus (Merkes et al., 2014), and secretions (Bylemans et al., 2017) of organisms. Previous studies have suggested that eDNA is mainly derived from fractions of cells or cellular organs, but can also be derived from fragmental DNA in water (Turner et al., 2014; Minamoto et al., 2016).

Many points regarding the general behavior of eDNA in water (Barnes and Turner, 2016) are still unclear, especially the state and degradation of eDNA in the water (Turner et al., 2015; Barnes and Turner, 2016). Rare and endangered species are thought to have a small population and release a small amount of DNA (Fukumoto et al. 2015; Sigsgaard et al., 2015; Pfleger et al. 2016; Doi et al., 2017; Sakata et al., 2017). Understanding the state and degradation of eDNA allows us to apply eDNA methods for various situations, for example, distribution evaluation for rare and endangered species with lower biomass/abundance. Even for organisms living in their known habitat, eDNA degradation could induce false negatives when their distributions are studied. For conservation surveys using eDNA, it is important to gather knowledge about eDNA degradation.

Many experiments have been conducted to reveal the detailed states and degradation rates of eDNA under various conditions (Thomsen et al., 2012; Barnes et al., 2014; Maruyama et al., 2014; Tsuji et al., 2017; Jo et al., 2019). In most cases, the eDNA degradation curves declined exponentially and quickly, often in less than a week (Thomsen et al., 2012; Barnes et al., 2014). Earlier meta-analyses for eDNA degradation (Collins et al., 2018) found that water conditions, such as salinity (Collins et al., 2018), water temperature (Tsuji et al., 2017; Jo et al., 2019), and pH (Barnes et al., 2014; Tsuji et al., 2017), influenced the eDNA degradation rate. In addition, the characteristics of DNA itself, such as its measured fragment length, affected the eDNA degradation rate (Bylemans et al., 2018, Jo et al. 2019). A general model for eDNA degradation can be applied to consider the eDNA state of species, including rare and endangered species, in their habitats. Therefore, we conducted a novel meta-analysis to model the effects of water conditions and DNA fragment length on the eDNA degradation rate. The previous meta-analysis used the half-life of the degradation curve as an index of degradation. To evaluate the eDNA degradation behavior, we expected that the degradation rate (slope of a simple exponential model) would be useful.

Our aim was to evaluate the effects of water conditions and DNA fragment length on the degradation rate by meta-analyses using previous published data. From the synthesis, we conducted a simulation to predict the maximum degradation rate in combination with water temperature and fragment length by modeling the quantile model.

## Methods

### 2.1. Search strategy

A Google Scholar search on September 9, 2020, using the search terms described below, returned 11,300 hits. The initial filtering of the articles was based on their abstracts: any articles that obviously had no relevance to eDNA degradation were discarded. After title screening, 1,000 articles remained. After abstract screening, 42 articles remained. We manually inspected these remaining articles and selected papers describing the degradation rate of eDNA using experiments or field settings (Table S1). We finally obtained the data from 28 articles (Tables 1 and S1) for the meta-analysis.

**Table 1.**
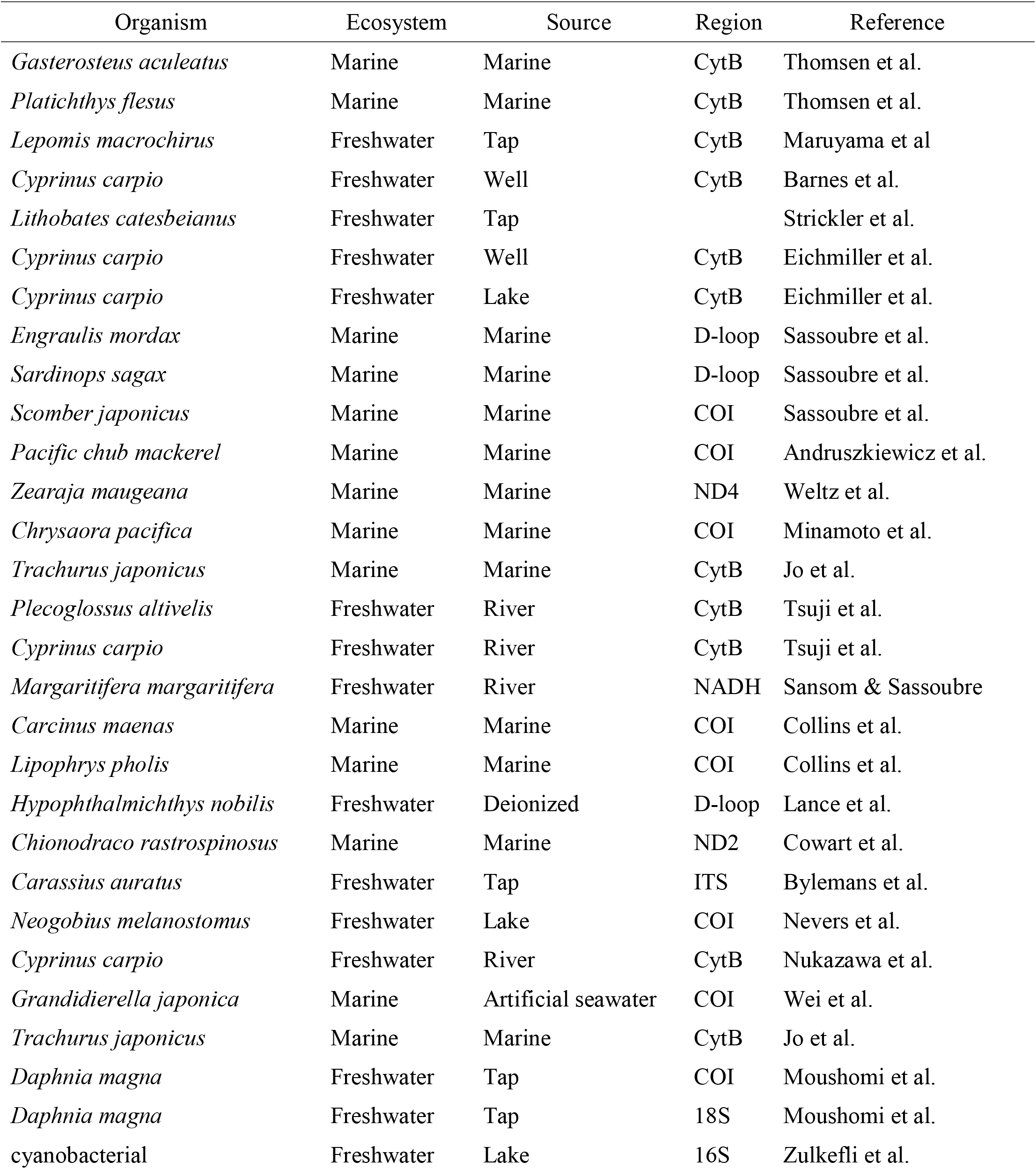
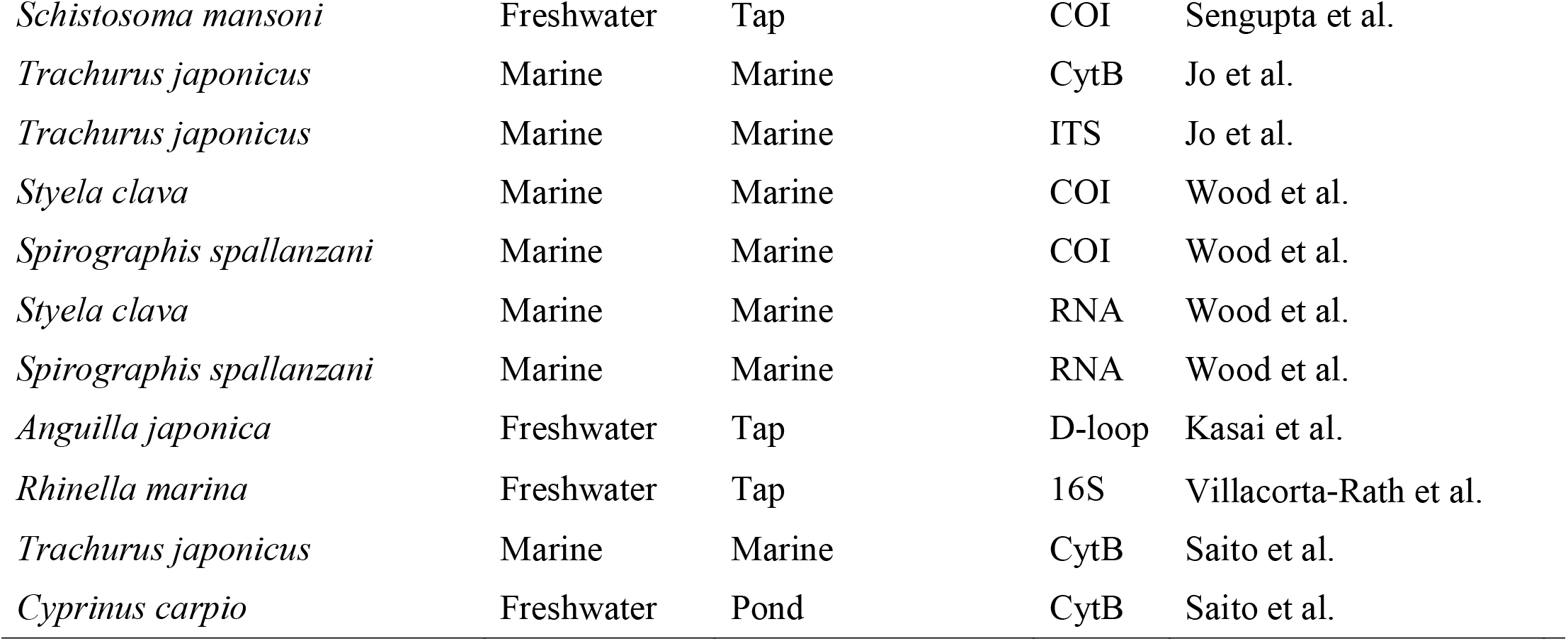
The organisms, ecosystem types (Ecosystem), water source (Source), and PCR-amplified DNA regions by quantitative PCR (Region) for all papers analyzed in this meta-analysis.

### 2.2. Data extraction

From the selected publications, we assembled a list of factors for eDNA degradation (Table S1). We collected the following factors and categories: “Ecosystem” was divided into marine and freshwater. “Source” was categorized into water sources (Freshwater: river, lake, well water, pond, tap water and deionized water; Marine: marine and artificial seawater). “Temperature” and “pH” refer to the water temperature and pH of the water sample for each experiment, respectively. “Region” and “Fragment length” refer to the amplified DNA region used for quantitative PCR (qPCR) and the number of amplified-DNA bases (bp), respectively.

We extracted the simple exponential slope (hereafter referred to as “degradation rate”) according to the simple exponential equation in each experiment:

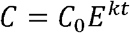

where C_0_ is the eDNA concentration at time 0, i.e., the initial eDNA concentration, and k is the degradation slope (rate) constant per hour. We used the standardized degradation rate per hour.

### 2.3. Statistical analysis and simulation

We performed the statistical analysis and graphics using R ver. 4.0.2 (R Core Team, 2020). We tested the differences in the eDNA degradation rate in measured DNA regions and water resources using a linear mixed-effect model (LMM) using “lme4” ver. 1.1.23 package with “lmerTest” ver. 3.1.2 package in R. We set each study as a random effect. We performed quantile models (QM) for 0.1, 0.5, and 0.95 quantiles for the regression. We employed the Bayesian mixed-effect quantile model using the “lqmm” function of “lqmm” package ver. 1.5.5 in R. In the QM, we set water temperature and fragment length as explanatory effects and each study as the random effect. We performed the Nelder–Mead algorithm using 10000 MCMC permutations with the Gauss–Hermite quadrature approach. We set the statistical alpha as 0.05 for parameter evaluation. We did not find a significant interaction (p > 0.1) between water temperature and fragment length, so we used the model excluding the interaction, i.e., eDNA degradation rate = water temperature + fragment length. We evaluated the QM models using the Akaike information criteria (AIC) of the different quantile models.

We simulated the combined effects of water temperature and fragment length and the maximum degradation rate under these conditions, using the obtained 0.95-quantile QM. We generated 100,000 random values for the combination of water temperature (ranging in published values from −1 to 35 °C; see the results) and fragment length used for the experiments (ranging in published values from 70 to 719) using “runif function in R, which generates a random number from the Mersenne-Twister method. We used 100,000 random values to predict the eDNA degradation rate from the 0.95-quantile QM (see results).

## Results

### 3.1. Experiments

The number of obtained time points for the eDNA degradation data ranged from 3 to 25 (mean: 8.3, median: 8.0, Fig. S1). Details of the site are listed as water sources (Table 1); 21 marine sites included sources from 19 freshwater sites, which included 4 river sites, 1 pond site, 3 lake sites, 2 sites of well water, and 9 experiments with tap or deionized water, and 1 artificial seawater site. The temperature for the experiments ranged from −1 to 35 °C (mean: 19, median: 20, Fig. S1). The fragment length used for the experiments ranged from 70 to 719 bp (mean: 150, median: 131, Fig. S1), and the fragment regions used were mainly Cyt B or COI regions in mitochondrial DNA (Table 1).

### 3.2 Degradation rate

The degradation rate for the eDNA degradation data ranged from 0.0005 to 0.7010 (mean: 0.1317, median: 0.0440, Fig. S1). Differences in PCR regions did not affect the rate of DNA degradation, nor did differences in water sources (Fig. 1A, B). There were no significant differences among the water sources and PCR regions (LMM, t < 1.859, p > 0.07).

**Figure 1.**
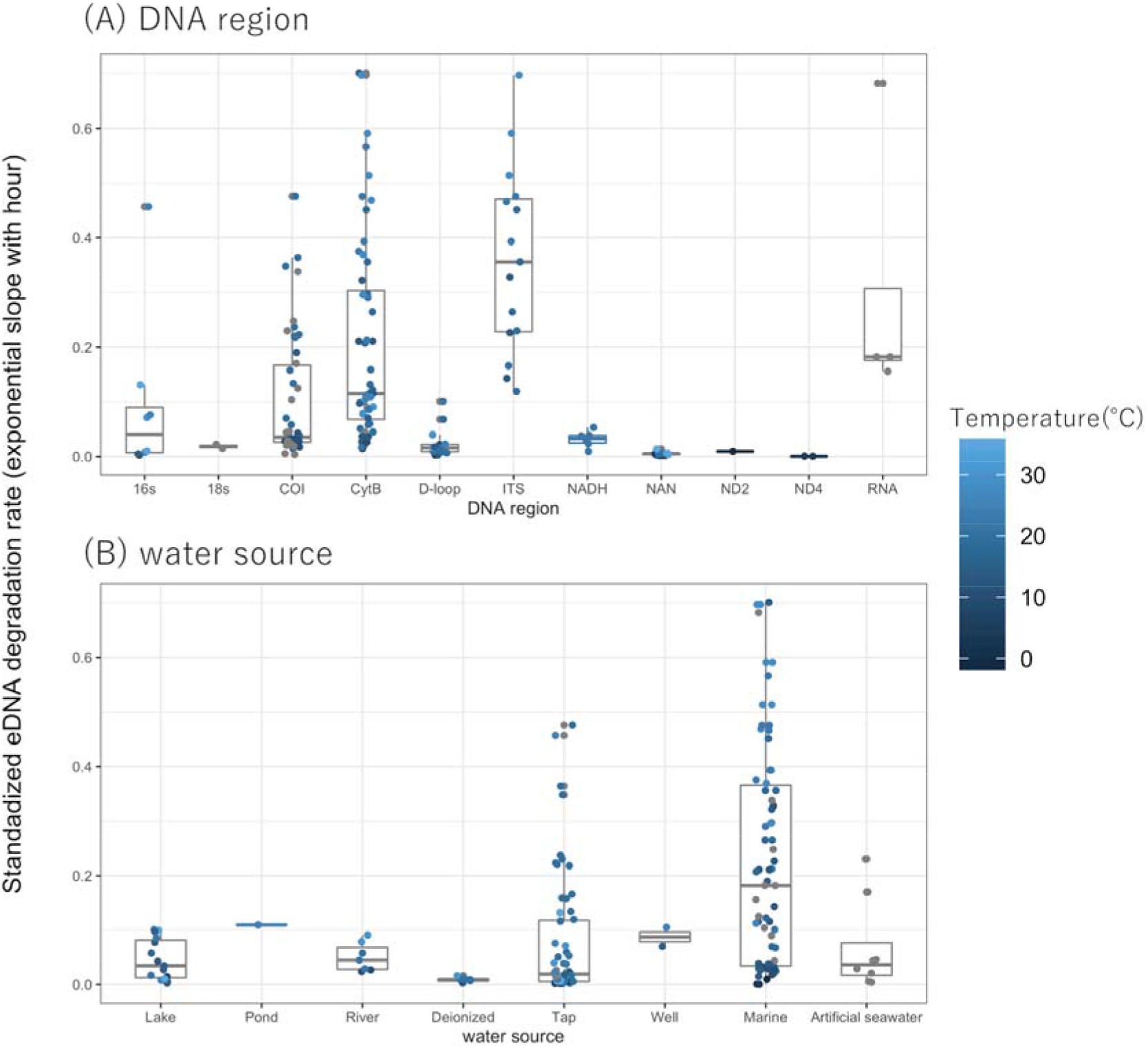
The eDNA degradation rate (simple exponential slope) with **(A)** DNA region and **(B)** water source. The dots indicate the individual eDNA degradation rate in each experiment in different ecosystems: seawater, blue; freshwater, red. The boxes and bars in the box plot indicate median ± inter-quartiles and ±1.5 × inter-quartiles, respectively.

### 3.3. Quantile model for temperature and fragment length

The relationship between eDNA degradation rate and water temperature showed that higher water temperatures accelerated eDNA degradation (Fig. 2A). Upon comparing the QM of 0.1-, 0.5-, and 0.95-quantiles, the QM with 0.95-quantile was observed to have the lowest AIC value (0.1-quantile:86.97, 0.5-quantile: −102.07, and 0.95-quantile: −208.47). Therefore, we simulated and discussed these data using the QM with a 0.95-quantile with a positive slope (slope = 0.020, Fig. 2A). The relationship between eDNA degradation rate and fragment length showed that longer fragments accelerated eDNA degradation (Fig. 2B). Most of the amplification regions of PCR primers designed for detecting environmental DNA so far were 200 bp or less. For fragment length, as for water temperature, the QM with 0.95-quantile had the lowest AIC value (0.1-quantile: 163.1 (df = 4), 0.5-quantile: −100.6, and 0.95-quantile: −127.5). Therefore, we simulated and discussed these data using the QM with a 0.95-quantile with a positive slope (slope = 0.197).

**Figure 2.**
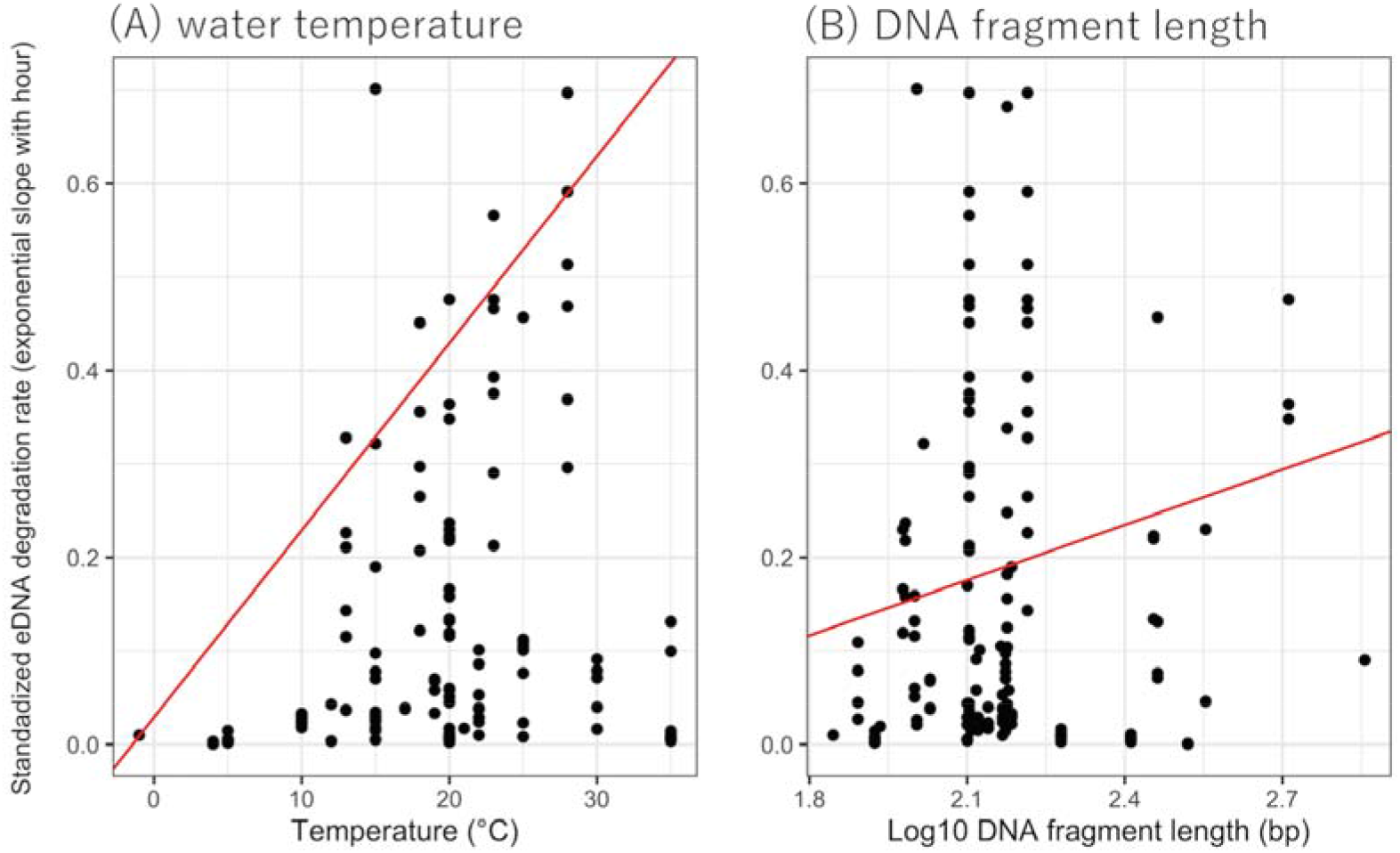
The relationship between standardized eDNA degradation rate per hour (simple exponential slope) with **(A)** water temperature and **(B)** DNA fragment length. The red lines show 0.95-quantile mixed-effect quantile models for each factor.

### 3.4. eDNA degradation simulation

Our QM simulation lead to plotting the eDNA degradation on a matrix of water temperature and fragment length (Fig. 3) and showed that the water temperature had a great influence on the eDNA degradation rate. At lower (e.g., 0 to 5 °C) and higher (e.g., 15 to 35 °C) water temperatures, we predicted that fragment length would have a smaller effect on the eDNA degradation rate, while at moderate (e.g., 5 to 15 °C) water temperatures, our prediction more clearly showed that the longer fragments would have a faster degradation rate. Thus, at moderate water temperatures, the fragment length should also be considered in evaluating eDNA degradation.

**Figure 3.**
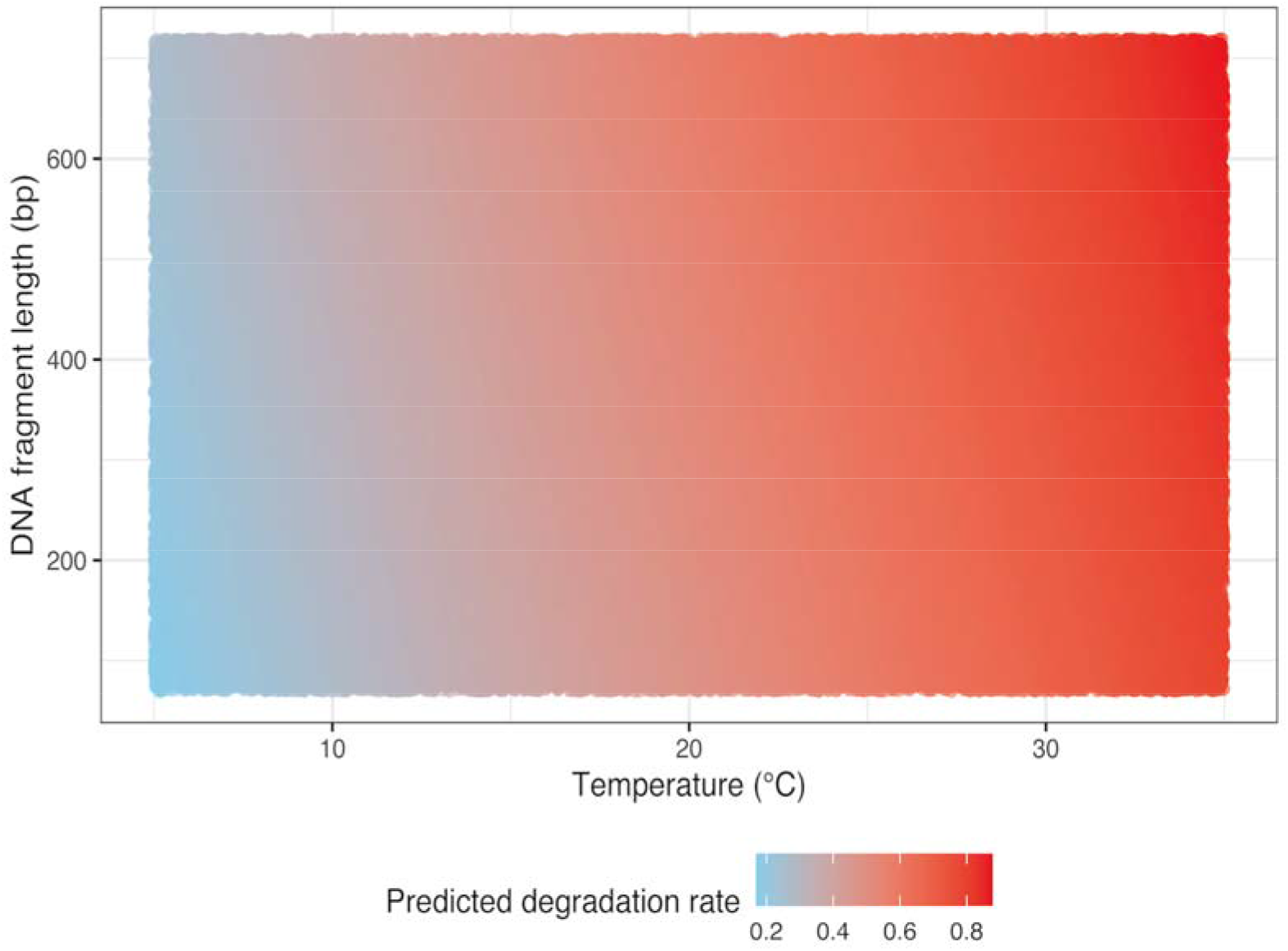
The simulation result for predicting eDNA degradation rate on the matrix of water temperature and fragment length.

## DISCUSSION

Our meta-analysis results showed that higher water temperatures and longer fragments accelerated eDNA degradation. These generally supported the effect of water temperature on the eDNA degradation rate in previous hypotheses for each condition and species (e. g., Strickler et al., 2015; Eichmiller et al., 2016; Lance et al., 2017; Tsuji et al., 2017; Jo et al., 2018; Kasai et al., 2020). Previous studies have assumed that water temperature does not directly affect eDNA degradation, but indirectly affects it through enzymatic hydrolysis by microbes and extracellular nucleases (Barnes and Turner 2016). At high temperatures, with increasing activity of microorganisms and extracellular enzymes, the eDNA in water would decompose more quickly (Barnes and Turner, 2016).

In addition, long DNA fragments were less likely to be detected than short fragments, and our meta-analysis supported the previous results. For example, Jo et al. (2017) suggested that the DNA degradation rate was higher in longer fragments (719 bp) than in shorter fragments (127 bp). Our simulation by QM indicated that shorter fragments were more likely to be retained when equivalent eDNA degradation occurred due to water temperature. When the eDNA degradation rates were very fast or very slow due to water temperature, the fragment length had a smaller effect on eDNA degradation than at other water temperature ranges. When the temperature-dependent degradation was very fast, we would expect that the eDNA would decompose regardless of the fragment length, probably because short fragments would be decomposed at a similar rate to longer ones, and hence, would not be retained. However, when temperature-dependent degradation occurs slowly, fragment length might influence the eDNA degradation rate because the longer DNA fragments could be retained, depending on their lengths. In our meta-analysis, we evaluated fragment lengths ranging from 70 to 719 bp, but there were no experiments in which longer fragments were measured. Recently, the long-PCR method has been developed for evaluating longer fragments of mitochondrial DNA (Deiner et al., 2017). By examining longer fragments of mitochondrial DNA, future studies can better understand the effect of fragment length on eDNA degradation.

There are some cases where eDNA has not been detected, even if the habitat of organisms has been confirmed. It has been pointed out that false negatives may involve eDNA degradation in the environment and eDNA measurement processes, such as water sample transport (Barnes and Turner 2016). By predicting the amount of eDNA degradation, we can estimate, for example, how much eDNA will be degraded by the time the water sample has been transported to the laboratory. If the amount of such degraded eDNA is not taken into consideration, species distribution and abundance/biomass may be underestimated, especially for low-density species such as rare and endangered species. Thus, we can apply the understanding and suppression of eDNA degradation to the detection of trace eDNA amounts. Similarly, we can apply the understanding of invasive distribution evaluation by eDNA because it is important to detect alien species in the early stages of invasion, when their abundance, i.e., eDNA concentration, may be low. Considering the rapid eDNA degradation in water, it is important to suppress any decomposition after obtaining the water sample. Several methods have been used to suppress eDNA degradation, including the addition of benzalkonium chloride (BAC) to transport water samples (Yamanaka et al., 2017; Takahara et al., 2020), ethanol or isopropanol fixations of water samples for transport (Doi et al. 2017), filtering at a water sampling site, and storing in RNAlater (Sigma-Aldrich, St. Louis, MO, USA) (Miya et al., 2016). These methods may be very useful for suppressing eDNA degradation, especially for environments with higher water temperatures and for detecting longer DNA fragments.

In conclusion, we found that higher water temperatures and longer DNA fragments generally accelerated eDNA degradation. We predicted the combined effects of water temperature and fragment length on the maximum eDNA degradation rate. Our meta-analysis and simulation provided new insights for future eDNA studies. We should note the limitations: The number of papers used for our meta-analysis was limited to 28 studies, and the data was limited especially for other environmental factors, such as UV, pH, and salinity, which are important factors for eDNA degradation (Mächler et al., 2018; Barnes et al., 2014; Tsuji et al., 2017; Lance et al., 2018; Collins et al., 2018). When data such as UV, pH, and salinity are obtained in addition to water temperature, more complex phenomena can be evaluated to determine the eDNA degradation rate in water. A greater understanding and accumulation of eDNA degradation data would improve future eDNA methods.

## Author Contributions

TS and HD designed the study; TS collected the data; TS and HD analyzed the data and interpreted the results; and TS and HD wrote the manuscript.

## Acknowledgments

We thank the authors of all the papers used for the meta-analysis. This study was supported by the Environment Research and Technology Development Fund (4-2004).

## Supplementary Material

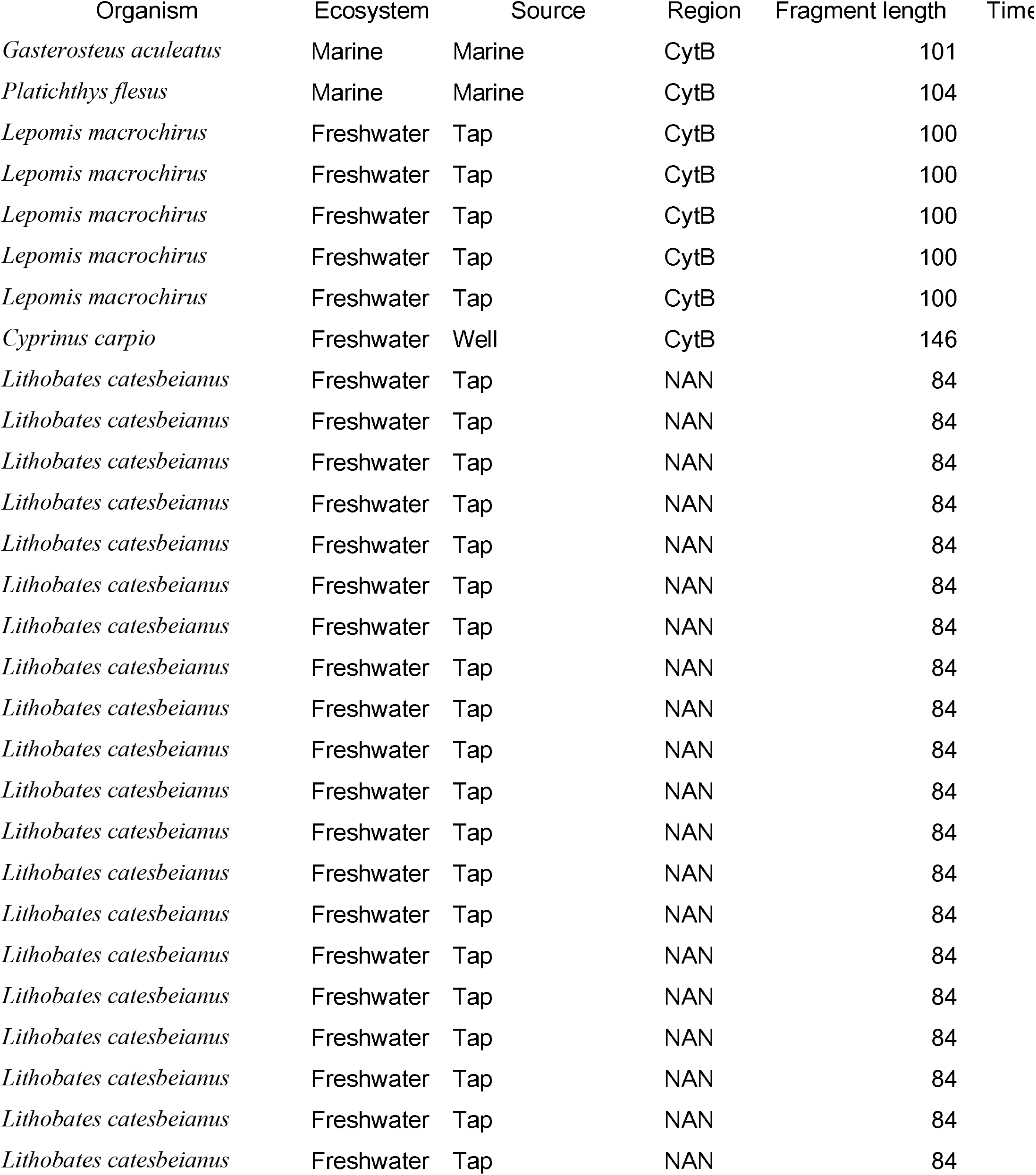
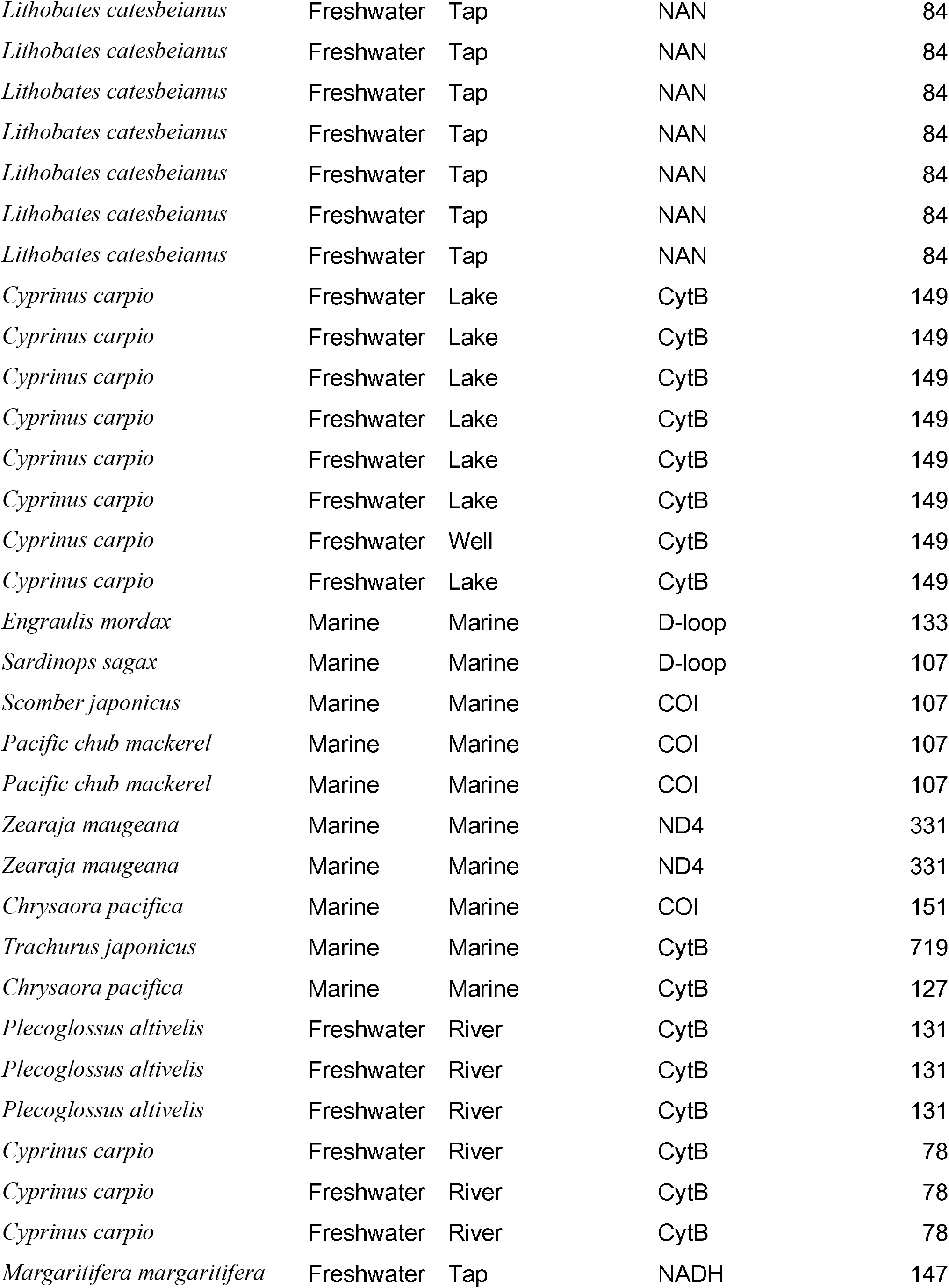
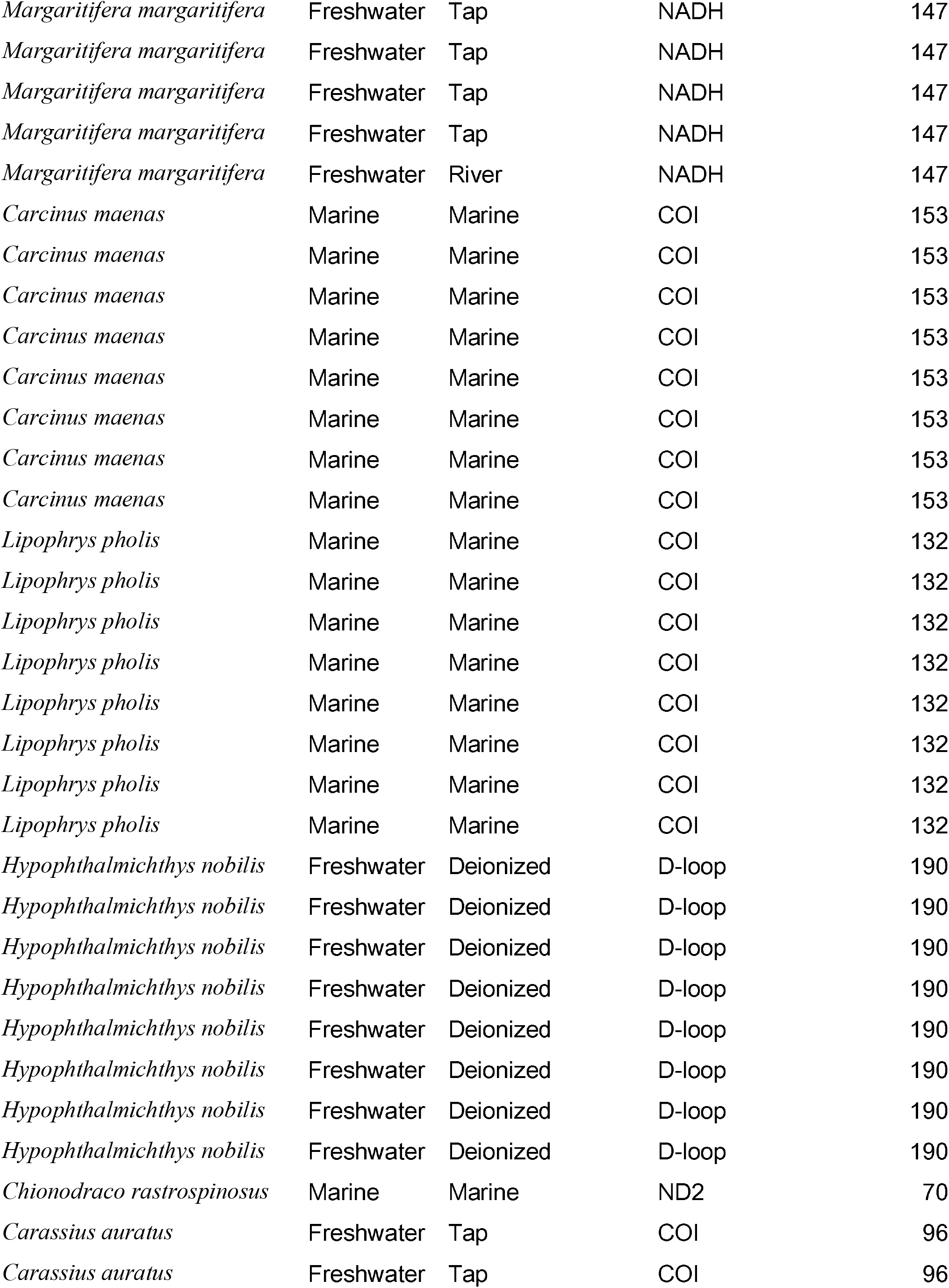
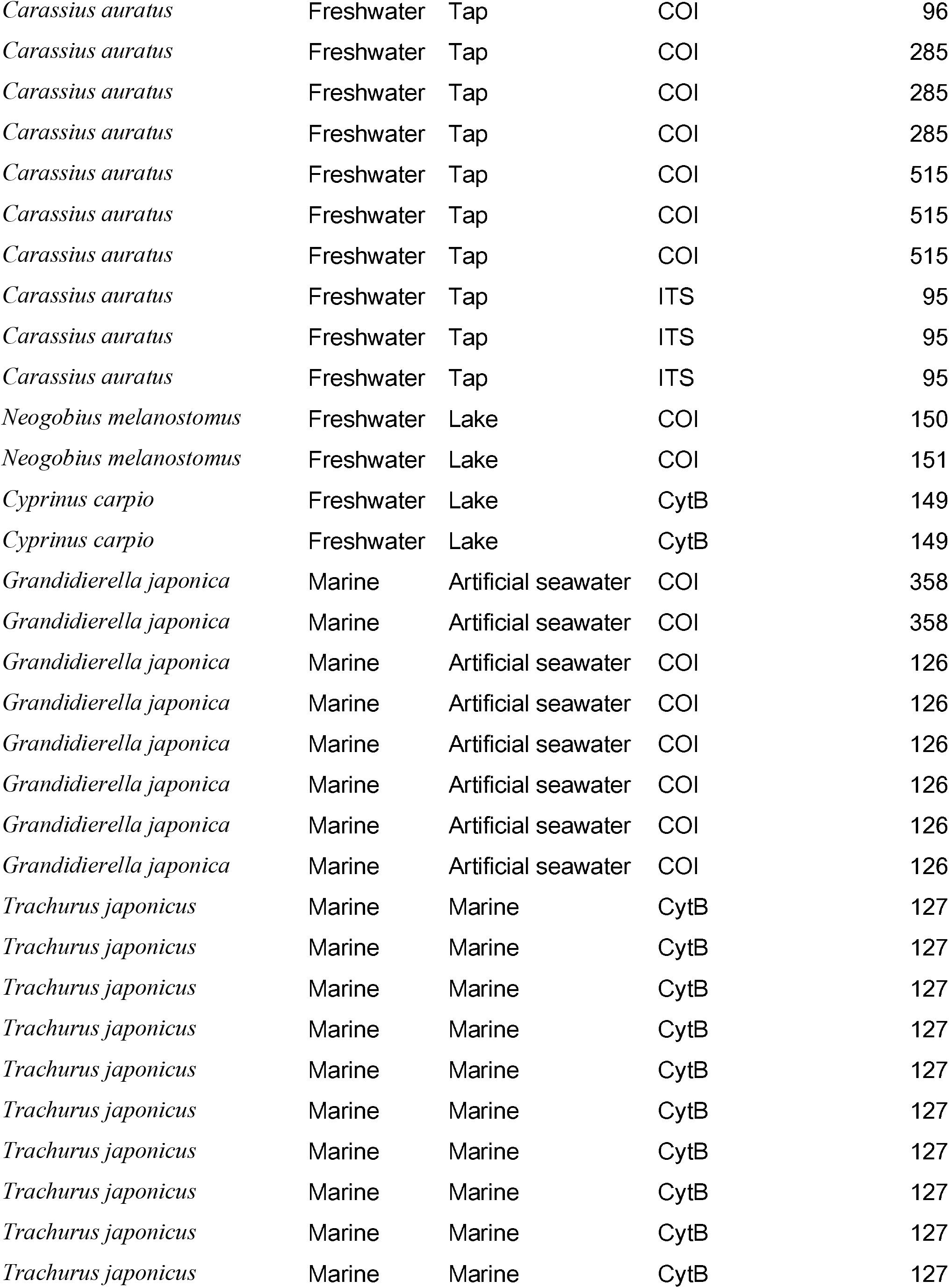
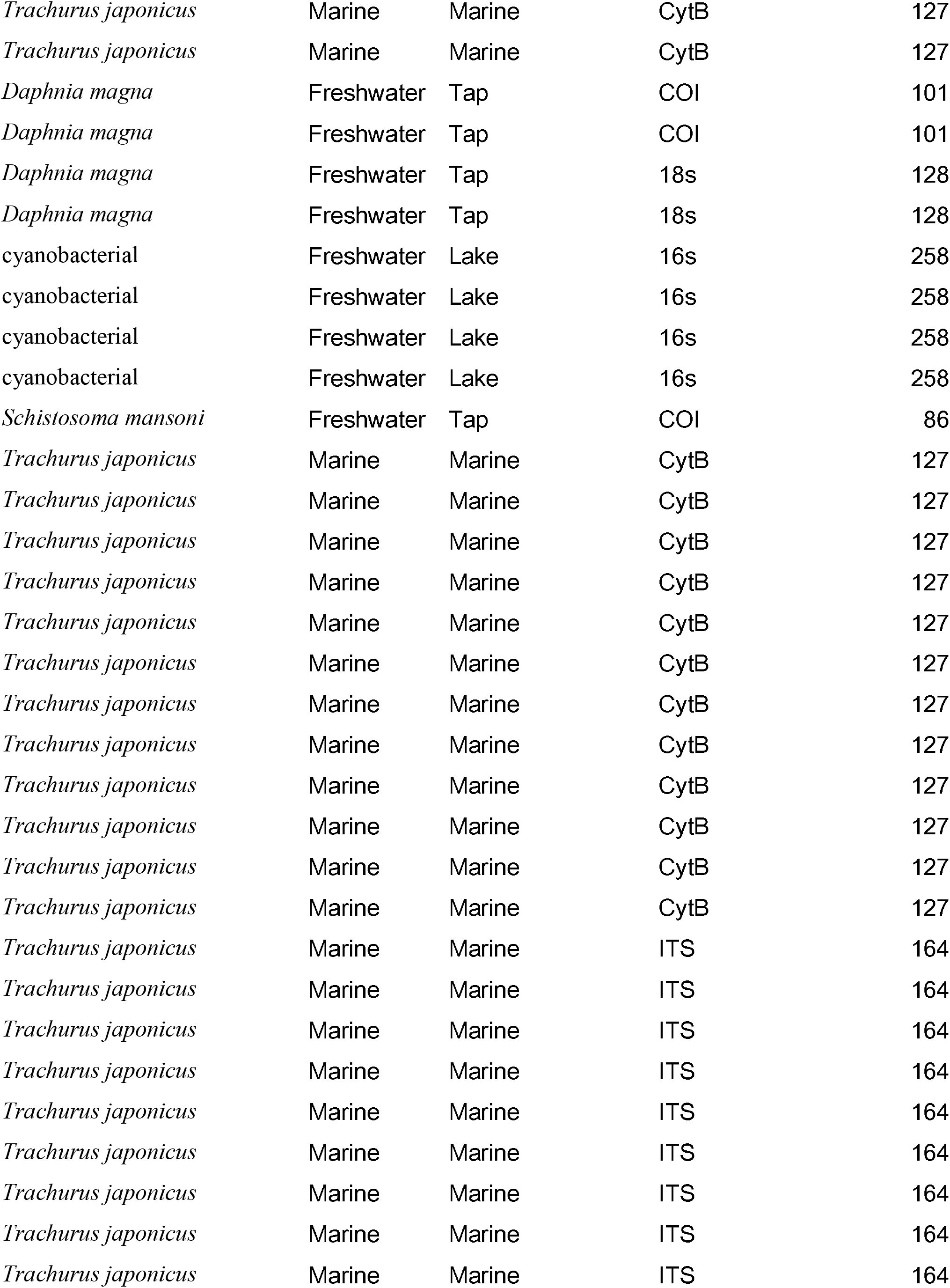
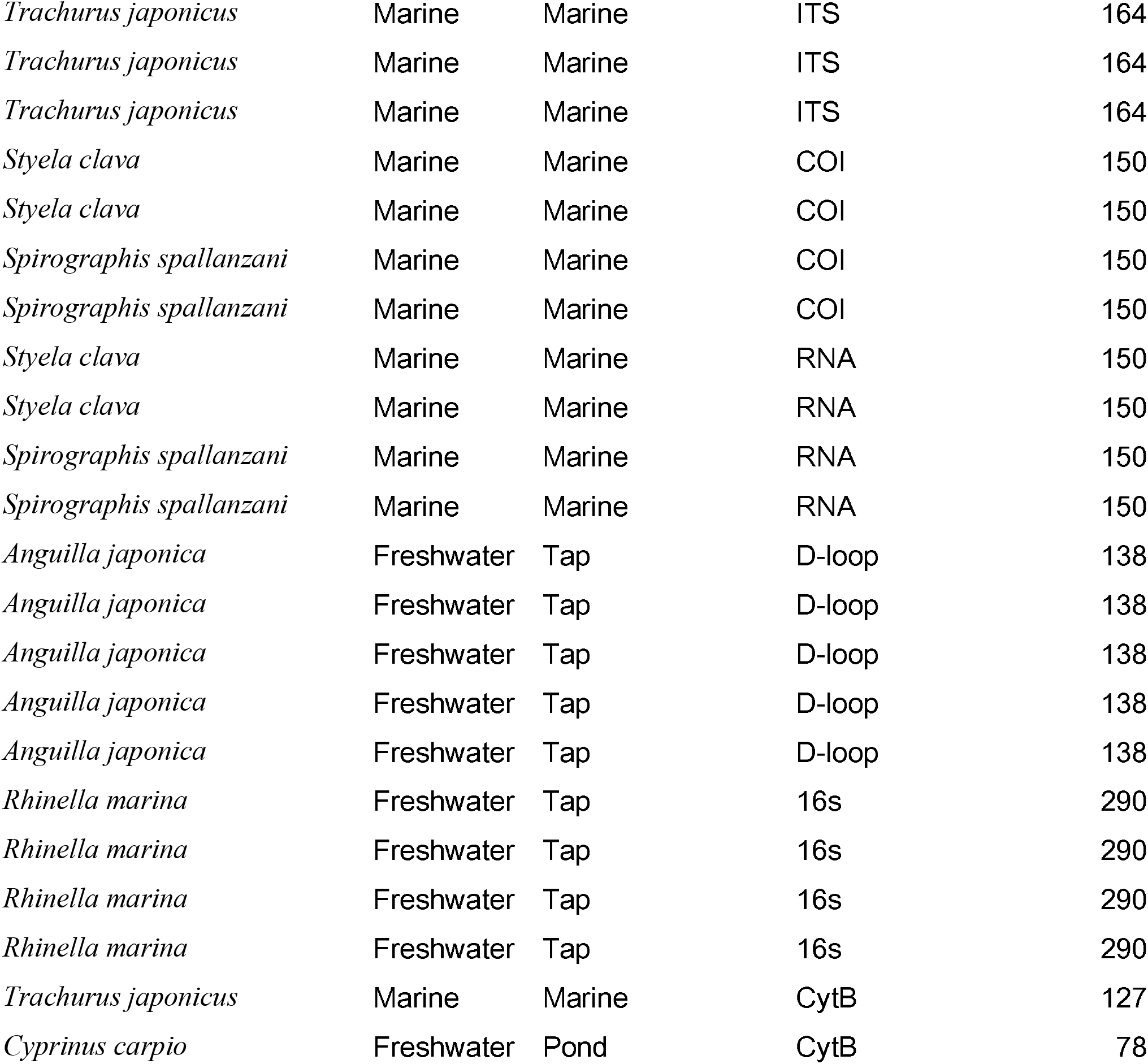

